# Effects of light / dark diel cycles on the photoorganoheterotrophic metabolism of *Rhodopseudomonas palustris* for differential electron allocation to PHAs and H_2_

**DOI:** 10.1101/2020.08.19.258533

**Authors:** Marta Cerruti, Heleen T. Ouboter, Viktor Chasna, Mark C. M. van Loosdrecht, Cristian Picioreanu, David G. Weissbrodt

## Abstract

Light/dark cycles can impact the electron distribution in *Rhodopseudomonas palustris*, a hyperversatile photoorganoheterotrophic purple non-sulfur bacterium (PNSB). Dynamic conditions during diel cycles are important for the physiology of PNSB, but the coupling between illumination patterns and redox balancing has not been extensively studied. For survival and growth, *Rhodopseudomonas* has developed different mechanisms to allocate electrons under dynamic growth conditions. Products such as hydrogen and poly-β-hydroxyalkanoates (PHAs) can form alternative electron sinks. A continuous culture, fed with a balanced nutrients medium, was exposed to three different conditions: 24 h continuous infrared illumination, 16h light/8h dark, and 8h light/16h dark. Light and dark phase durations in a cycle determined the energy availability level (light) and the attainment of a stationary state. Under long dark phases, the acetate substrate accumulated to levels that could not be depleted by growth in the light. Under short dark phases, acetate was rapidly consumed in the light with most of the phototrophic growth occurring under acetate-limiting conditions. Under diel cycles, substrate uptake and growth were unbalanced and *Rhodopseudomonas* shunted the excess of carbon and electron flow first toward PHAs production. Only secondarily, when PHA storage got saturated, the electron excess was redirected toward H_2_. A numerical model described well the dynamics of biomass and nutrients during the different light/dark cycle regimes. The model simulations allowed determination of stoichiometric and kinetic parameters for conversion by *Rhodopseudomonas*. Understanding the inherent process dynamics of diel light cycles in purple sulfur bacteria cultures would enable optimization procedures for targeted bioproduct formation.

**Importance:** Purple non-sulfur bacteria (PNSB) are important anoxygenic phototrophic microorganisms that take part in numerous environmental processes, based on their metabolic versatility. *Rhodopseudomonas palustris* is a model photosynthetic bacterium of the PNSB guild. Light cycles influence deeply its physiology. Poly-β-hydroxyalkanoates (PHAs) and biohydrogen are two of the most studied metabolic products of *Rhodopseudomonas*, because of their biotechnology potential besides involvement in carbon and electron allocations in its metabolism. Their production mechanisms have often been described as competitive, but the rationale behind the production of one or the other compound has not been elucidated. Here, we found that under light / dark cycles an excess of organic substrate was first directed toward PHAs production, and only when this pathway was saturated H_2_ was produced. Understanding the dynamics of carbon and electron allocation under intermittent light cycles enhances our knowledge on PNSB metabolisms and paves ways to manage the formation of targeted bioproducts.

## Introduction

Purple non-sulfur phototrophic bacteria (PNSB) form a guild of hyper-versatile anoxygenic phototrophs (1), able to grow on different organic and inorganic substrates (2). *Rhodopseudomonas palustris* is one model PNSB (3). It can produce compounds of industrial interest, such as single cell proteins (4), carotenoids (5), hydrogen gas (H_2_; used as biofuel) and poly-β-hydroxyalkanoates (PHAs; used as bioplastics) (6,7). Under light and in presence of organic carbon, *Rhodopseudomonas* grows photoorganoheterotrophically even in absence of external electron acceptors. Under dark, growth is possible on sugars with external electron acceptors (as nitrate, trimethylamine-N-oxide or dimethyl sulfoxide) (8,9).

Light is crucial for phototrophy by providing energy to cells. PNSB capture the photonic energy and couple the light-driven oxidation of the photopigments to an electron transfer through membrane bound-enzymes. The trans-membrane gradient of hydrogen protons (H^+^) supports ATP synthesis through cyclic photophosphorylation (10). NADH is mainly produced during catabolic processes. In some cases, a reverse electron flow takes places, and the NADH-dehydrogenase catalyze the proton transfer from the ubiquinone pool to NAD^+^. The control of redox balance is crucial for cell survival and growth. The ratio between NADH/NAD^+^ is important for intracellular redox homeostasis (11). PNSB generate NADH in three different ways (Figure 1), namely: 1) anabolic processes (12), 2) light driven reactions through the quinone pool (1); 3) reverse electron transfer (13).

**Figure 1.**
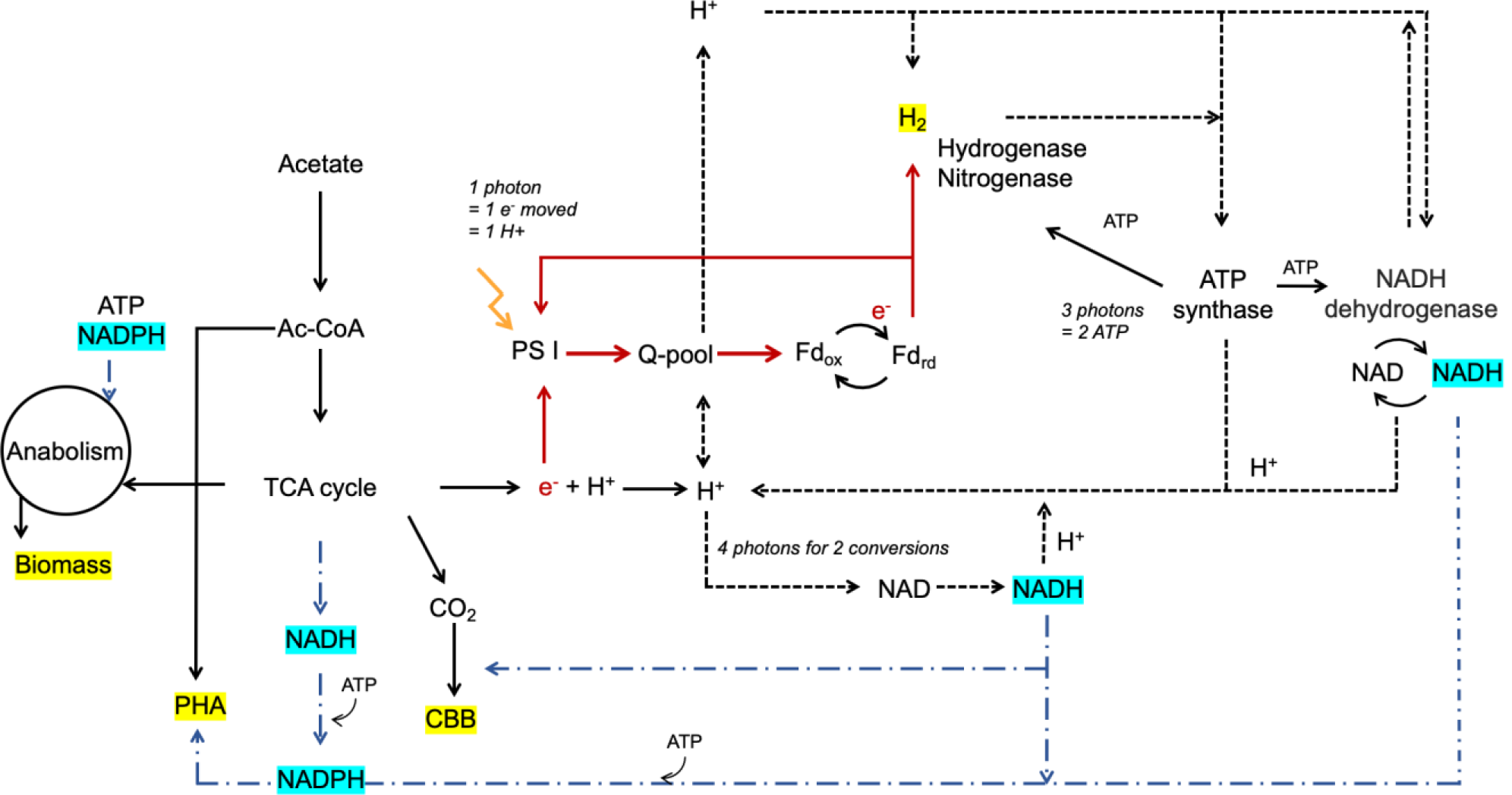
Schematic representation of the reducing power allocation in *Rhodopseudomonas*. <u>Red lines</u>: electron flow; <u>Black dashed lines</u>: proton transfer; <u>Blue dotted lines</u>: NADH is normally produced in the catabolic processes, whereas NADPH is used in the anabolic processes, such as PHAs formation. Biomass and CBB are the primary electron sinks in PNSB. PHAs and H_2_ are two of the other possible electron sinks.

To cope with possible redox imbalances, mainly arising in anabolic processes, PNSB redistribute the electrons toward different routes (14). In *Rhodopseudomonas*, the first and most important electron sink is the biomass itself, that receives the intermediates of the tricarboxylic acid cycle and the NADPH produced there (15). Secondly, the Calvin-Benson-Bassham (CBB) cycle is constitutively active also under photoorganoheterotrophy as central recycling mechanism of redox cofactors coming from the anabolic process (15,16). Under photoorganoheterotrophic anaerobic conditions, a surplus of electrons can be disposed through PHA or H_2_ production. Both electron sinks are normally produced under nitrogen limitation. Nitrogenase-mediated H_2_ production has been extensively described (1). H_2_ is a side-product of the nitrogen fixation process, that can only happen when no preferred nitrogen source like ammonium or glutamate is available (17–19). Hydrogenase-mediated H_2_ production has been reported in presence of an oxidized organic compound (*i.e*. malate) (21). Several bacterial species accumulate PHAs in the cells as a mean of carbon and electrons storage under normal conditions (22). In lab-scale experiments, to achieve an over production, these storage polymers are synthetized when acetyl-CoA and NADPH are in excess but nutrients like nitrogen, sulfur or phosphorus are limiting (23). Similar to other bacteria, PNSB can produce PHAs (24) as a mean of carbon and electron balance, utilizing the reducing power to build the storage polymers (25,26). PHAs production in PNSB occurs under balanced growth, and it increases under nutrient limitations (27). The two processes of H_2_ and PHA production are considered competitive electron dissipation pathways (28,29). However, the hyper-versatility of PNSB metabolisms has not enabled to identify an univocal response to redox imbalances.

In natural environments, phototrophs are subjected to diel (*i.e*., 24-h period) light / dark cycles. Since light is responsible for the production of energy and reducing power, the irradiation patterns impacts the cellular physiology. The light and dark cycles impact cells redox and ATP balance, subjecting cells to metabolic switches. How it affects the internal electron allocation patters and how PNSB respond to these switches remains puzzling. Allocation of reducing power toward the aforementioned routes is known, but mechanisms that govern the preferential electron flow have not been explained.

Here, we aimed to elucidate the effects of diel light / dark cycles on the physiology and electron allocation in *Rhodopseudomonas palustris*, isolated from an in-house PNSB enrichment culture for nutrient removal from wastewater (30). The dynamic conditions were hypothesized to influence the physiology of purple bacteria, as the energy source is intermittently provided. The process mimics a potential natural scenario, where day and night cycles are applied and create a redox imbalance. We elucidated the preferential carbon and electron redistribution toward the most important competitive pathways of PHAs and H_2_ formation during diel light / dark cycles.

## Results

Three regimes for continuous cultivation of *Rhodopseudomonas palustris* were tested: continuous illumination, cyclic 16 h light / 8 h dark, and cyclic 8 h light / 16 h dark. The biomass and CO_2_ yields (*Y*_*XS*_ and *Y*_*CX*_) and the maximum biomass-specific growth rate (μ_*max*_) were estimated using data from the 16 h light / 8 h dark experiment (Table 1). The calibrated model was used to predict the behavior of the 8 h light / 16 h dark cycles.

**Table 1.**
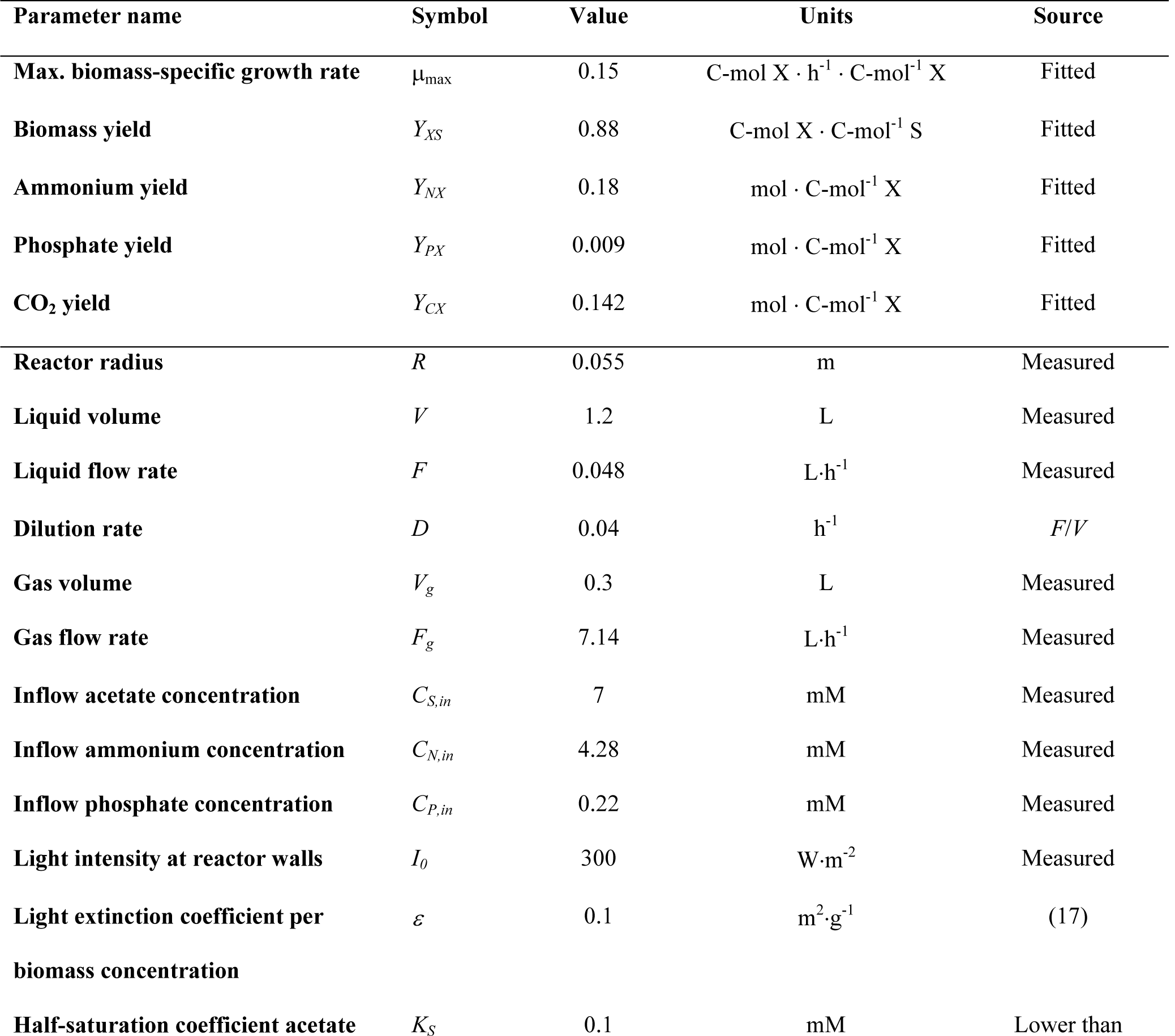
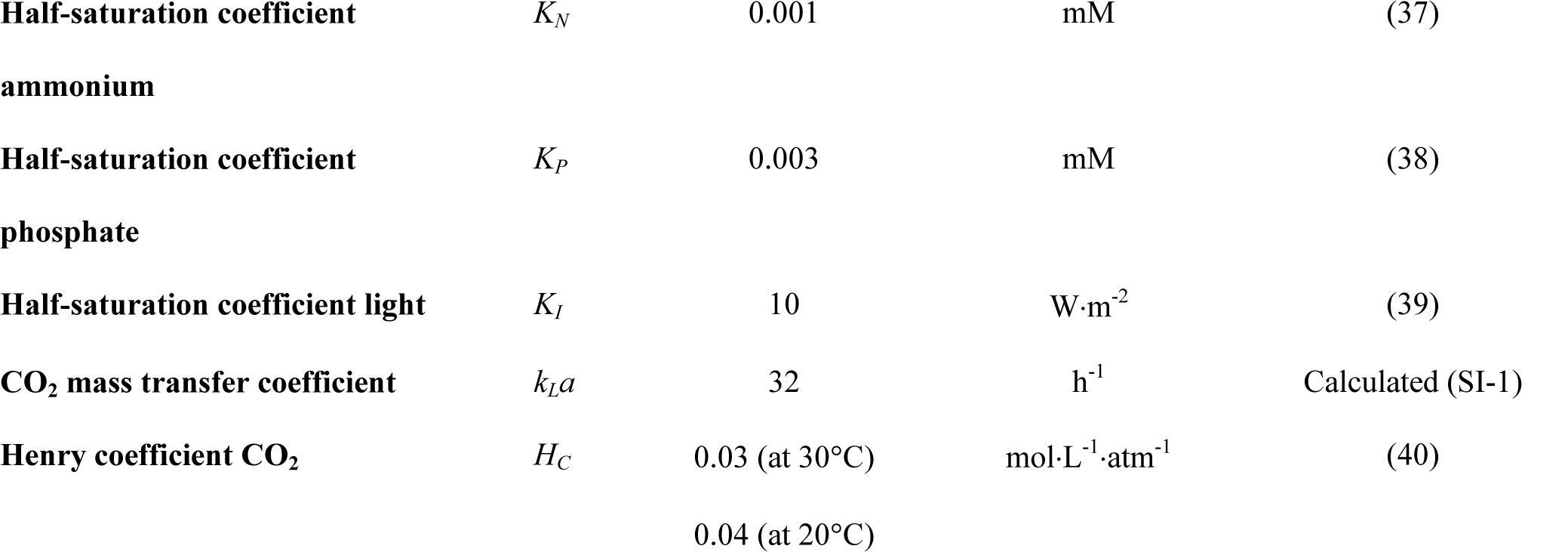
Model parameters. The maximum biomass-specific growth rate and yields were fitted to experimental data, while the other parameters were measured during experiments, calculated or obtained from literature.

| Parameter name | Symbol | Value | Units | Source |
| --- | --- | --- | --- | --- |
| Max. biomass-specific growth rate | $\mu_{\max}$ | 0.15 | $\text{C-mol X} \cdot \text{h}^{-1} \cdot \text{C-mol}^{-1} \text{ X}$ | Fitted |
| Biomass yield | $Y_{XS}$ | 0.88 | $\text{C-mol X} \cdot \text{C-mol}^{-1} \text{ S}$ | Fitted |
| Ammonium yield | $Y_{NX}$ | 0.18 | $\text{mol} \cdot \text{C-mol}^{-1} \text{ X}$ | Fitted |
| Phosphate yield | $Y_{PX}$ | 0.009 | $\text{mol} \cdot \text{C-mol}^{-1} \text{ X}$ | Fitted |
| CO<sub>2</sub> yield | $Y_{CX}$ | 0.142 | $\text{mol} \cdot \text{C-mol}^{-1} \text{ X}$ | Fitted |
| Reactor radius | $R$ | 0.055 | m | Measured |
| Liquid volume | $V$ | 1.2 | L | Measured |
| Liquid flow rate | $F$ | 0.048 | $\text{L} \cdot \text{h}^{-1}$ | Measured |
| Dilution rate | $D$ | 0.04 | $\text{h}^{-1}$ | $F/V$ |
| Gas volume | $V_g$ | 0.3 | L | Measured |
| Gas flow rate | $F_g$ | 7.14 | $\text{L} \cdot \text{h}^{-1}$ | Measured |
| Inflow acetate concentration | $C_{S,in}$ | 7 | mM | Measured |
| Inflow ammonium concentration | $C_{N,in}$ | 4.28 | mM | Measured |
| Inflow phosphate concentration | $C_{P,in}$ | 0.22 | mM | Measured |
| Light intensity at reactor walls | $I_0$ | 300 | $\text{W} \cdot \text{m}^{-2}$ | Measured |
| Light extinction coefficient per biomass concentration | $\varepsilon$ | 0.1 | $\text{m}^2 \cdot \text{g}^{-1}$ | (17) |
| Half-saturation coefficient acetate | $K_S$ | 0.1 | mM | Lower than |

### Biomass production of *Rhodopseudomonas palustris*

Under continuous illumination, the biomass reached a constant concentration of 12.10 ± 0.44 C-mmol L^-1^, corresponding to 88% of carbon or 99% of reduction equivalents (COD) provided in the influent. Around 11% of the carbon present in the inflow was recovered as CO_2_ (0.112 C-mmol L^-1^) in the off-gas. HPLC results showed no residual acetate in the bulk liquid at steady state.

Under cyclic operations (Figure 2), the system reached a stable behavior with similar changes in concentrations during each cycle. Within a cycle a clear dynamic pattern was observed. Under dark, the biomass did not grow and was only washed-out. The biomass wash-out rate (*F*/*V*·*C*_*X*_) was very well represented by the numerical model (Figure 2A,B). The corresponding change in acetate, ammonium and phosphate concentrations during the dark phases matched with the expected change based on the feeding rate. No conversion occurred in the dark.

**Figure 2.**
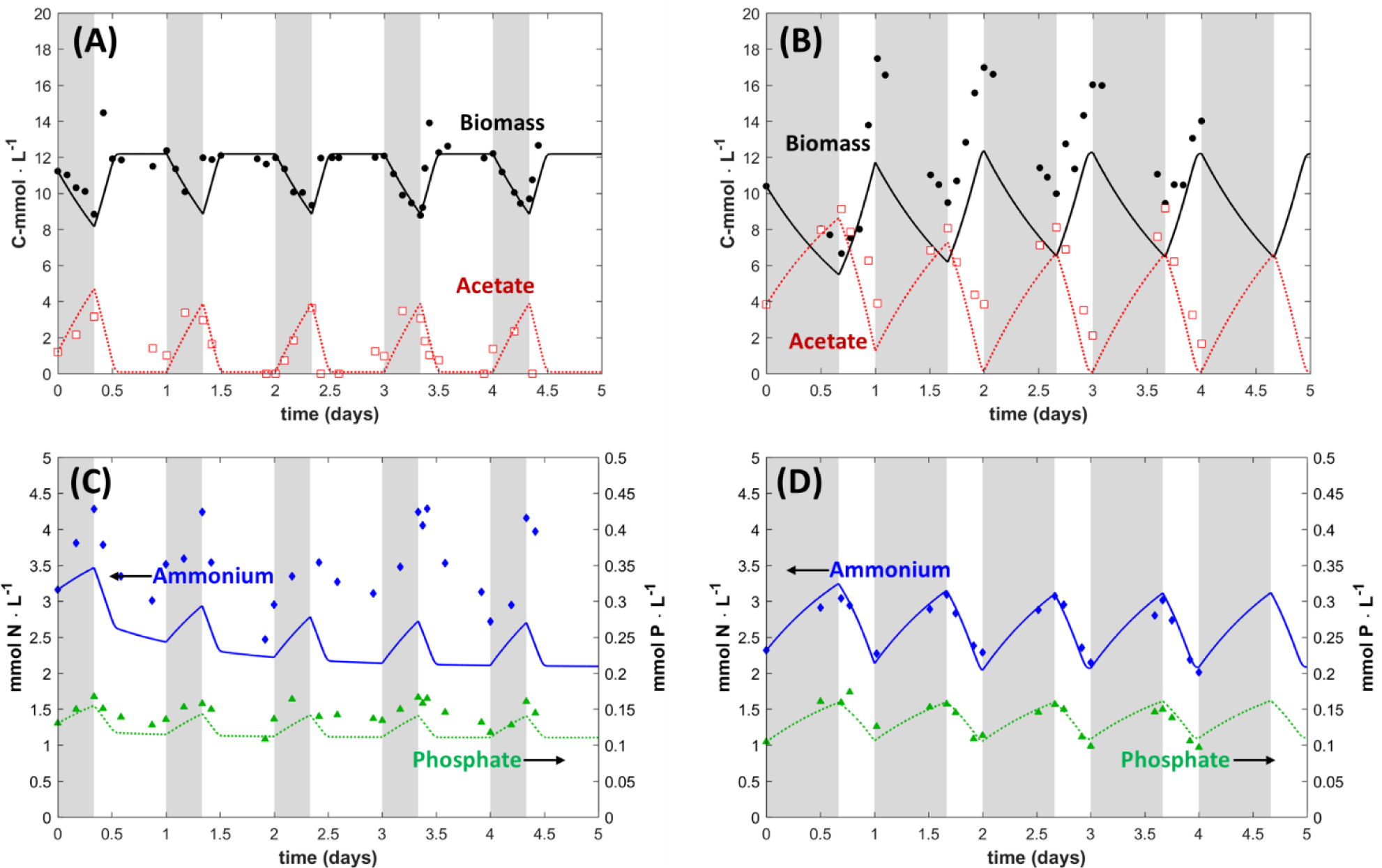
Dynamics of biomass, acetate, ammonium and phosphate concentrations during light-dark cycles: (A),(C) 16 h light / 8 h dark; (B),(D) 8 h light / 16 h dark. Gray areas represent the dark periods. (A),(B) biomass (black circles) and acetate (red open squares) measured concentrations, with lines being the model results. The biomass concentration achieved a stationary state, while acetate reached very low (limiting) concentrations after 2 h of the 16 h light periods. The biomass increased without reaching a steady state during the 8 h light periods, while acetate was not fully consumed. (C),(D) ammonium (blue diamonds) and phosphate (green triangles) measured concentrations, with lines being the model results. Concentrations of N and P were not limiting for the system in any illumination phase.

Under 16-h light, the biomass accumulation exceeded the dilution rate *D*=*F*/*V*. An initial growth rate (assumed to be µ_max_) of 0.15 h^-1^ was observed, and reached a steady value (µ = *D* = 0.04 h^-1^) after 2 h exposed to light (Figure 2A). The acetate concentration reached a minimal steady value when the biomass growth rate achieved the steady state.

Under shorter 8-h light periods, the biomass growth rate did not reach a stationary-state in a cycle, being around 0.1 h^-1^. Acetate was not depleted at the end of the light phase (Figure 2B). The biomass increase changed from 11 C-mmol L^-1^ on first cycle to 5 C-mmol L^-1^ on fourth cycle. The growth rate µ during the 8 h light period was approximately 0.12 h^-1^ thus lower than the µ_max_. The estimated biomass yield on acetate in the light was 0.88 C-mol X C-mol^-1^ acetate for both the 16 h light and the 8 h light phases, equal to the yield in continuous light regime.

### Carbon, nitrogen and phosphorus assimilation

Under continuous illumination, no residual acetate was detected in the bulk liquid. Under 16 h light, acetate was almost fully depleted, down to a concentration around 1 C-mmol L^-1^, as also very well represented by the numerical model. Under 8 h light, acetate was still present in the bulk liquid in concentrations ranging from 4 to 1.5 C-mmol L^-1^. In this case, the model overestimates the rate of acetate consumption, *i.e*. predicting acetate depletion at the end of the light phase.

Nitrogen and phosphorus sources were provided in excess, and never became limiting. Under light, N and P were consumed, resulting in a decrease in their concentrations (Figure 2C,D). Under dark, the fed N and P accumulated since not assimilated in biomass. At the end of the dark phase, 3.6 ± 0.6 N-mmol L^-1^ and 0.14 ± 0.01 P-mmol L^-1^ remained the bulk liquid. N and P dynamics showed irregular behaviour under 16 h light (Figure 2C). Therefore, the *Y*_*NX*_ and *Y*_*PX*_ were fitted on the 8 h light experiment (Figure 2D). The determined yields (*Y*_*NX*_ = 0.18, *Y*_*PX*_ = 0.009) were very close to the theoretical yields resulting from considering the biomass elemental composition (*Y*_*NX*_ = 0.18, *Y*_*PX*_ = 0.014) measured in (35). With these yields, the N and P dynamics during the 8 h light cycles were well represented by the model: accumulation under dark and consumption under light.

### CO_2_ production follows the biomass growth patterns

The CO_2_ production rate was constant at 0.075 ± 0.001 mmol h^-1^ during continuous illumination (Figure 3A): the chemostat achieved a stationary operation. Under light/dark cycles, CO_2_ emission in the off-gas decreased in the dark to 0.03 mmol h^-1^ (Figure 3B). Possibly, the registered baseline value can be due to the detection limit of the MS instrument. Under 16 h light, CO_2_ emission increased rapidly within the first hour, reaching 0.25 mmol h^-1^, following the biomass growth rate with acetate uptake. Once the acetate reached the minimal level, the CO_2_ production also decreased to a stable level of 0.07 mmol h^-1^ after the second hour of illumination, reflecting the steady state operation. The numerical model reproduced the observed trends during light phase, but the residual CO_2_ production in the dark phase was not accounted for.

**Figure 3.**
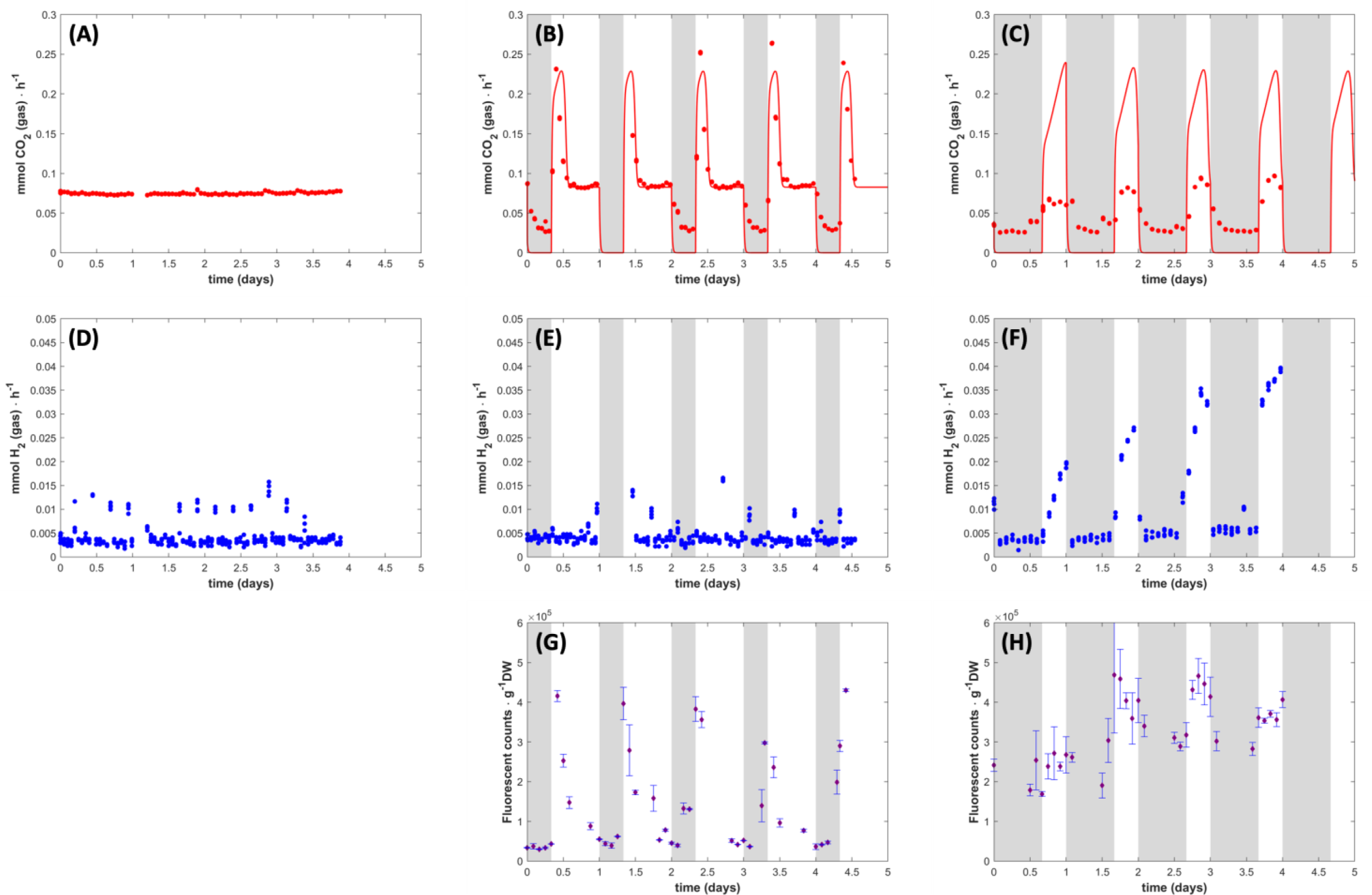
Dynamics of CO_2_, H_2_ and PHA during light-dark cycles: (A),(D) continuous illumination; (B),(E),(G) 16 h light / 8 h dark; (C),(F),(H) 8 h light / 16 h dark. Gray areas represent the dark periods. (A),(B),(C) CO_2_ production rate - measured (red circles) and computed (lines). CO_2_ production was constant during continuous illumination, but peaks appeared during light-dark cycles, correlated with the acetate uptake. (D),(E),(F) measured H_2_ production (blue circles). Constant low level H_2_ was produced during continuous illumination and 16 h light cycles, but H_2_ production strongly increased in each light period of the 8 h light cycles. (G),(H) PHAs fluorescent counts per gram biomass in the light/dark experiments. A PHAs peak was measured at the beginning of each of the 16 h light phases, however less clear pattern could be detected under 8 h light.

Under 8 h light / 16 h dar, the CO_2_ production did not achieve a steady state. In the light phase, the CO_2_ production increased with 0.05 to 0.08 mmol h^-1^ during the cycles, reaching a peak after 4 h (0.88 ± 0.013 mmol h^-1^), which was described only qualitatively by the model (Figure 3C). The reason for the discrepancy between model and experimental data is the larger computed acetate consumption rate, which leads to a higher CO_2_ formation rate in the model results. In both illumination regimes, the CO_2_ production dropped to the same stable value of 0.033 mmol h^-1^ during the dark phase.

### H_2_ is produced only during the short light periods

A very low H_2_ production was recorded during the continuous illumination with rates of around 0.004 mmol h^-1^ (Figure 3D). Occasionally, unexplained H_2_ spikes were recorded. During the 16 h light experiment, dark-phases hydrogen production was negligible and no specific production pattern was measured in the light (Figure 3E). The very low H_2_ concentration may relate to an instrumental offset. However, H_2_ was produced constantly during the 8 h light periods (Figure 3F), with an increase of 0.02 ± 0.01 mmol h^-1^ per cycle and reaching a maximum of 0.05 mmol h^-1^. The biomass-specific rate of H_2_ production under 8 h light was 10 times higher (0.156 mmol H_2_ h^-1^ g^-1^ DW) than under all other conditions (0.014 mmol H_2_ h^-1^ g^-1^ DW).

### PHA production follows CO_2_ formation under 16 h light but not under 8 h light

Due to the low biomass concentration, a traditional PHA extraction and GC quantification (41) was not possible. Fluorimetry was used to detect PHAs in the biomass, giving a relative quantification (34). PHAs were not detectable under continuous illumination. A fluorescence count peak was detectable after 2 h from the light switch in the 16-h light phases (3.8 ± 0.5·10^5^ fluorescent counts g^- 1^ DW) (Figure 3G), concomitantly to the peak of CO_2_ production. The PHAs decreased constantly after the peak, reaching a baseline at 3·10^4^ fluorescent counts g^-1^ DW. Under 8 h light / 16 h dark cycles (Figure 3H), PHA production during light was less pronounced. More PHAs were detectable in the dark compared to the 16 h light regime. These observations indicate that the diel light regimes induced PHA formation, with the long light regime generating a typical feast-famine behavior (42), while the short light regime stimulated a higher PHAs content in the cells.

## Discussion

### Acetate rather than light drives the metabolic responses of *Rhodopseudomonas*

Light energy source is central for the catabolic processes in PNSB. Organic substrates are used as C-source for biomass synthesis and as electron donors. When no external electron acceptor is present, the difference in degree of reduction between substrate and biomass has to be balanced with internal electron reallocation processes. The dynamics of the change in N and P concentrations were congruent with the biomass formation and their measured yields were close to the elemental biomass composition (*CH*_1.8_*O*_0.38_ *N*_0.18_*P*_0.014_) determined in another study (35). Carbon balances closed for the continuous illumination experiment and for the 16 h light / 8 h dark experiment, but not for the 8 h light / 16 h dark cycles, where the C-balances seemed to indicate a net production (Table 2). PHA granules that were always present in the cells under 8 h light / 16 h dark may have led to an absorbance overestimation of the biomass present in the reactor.

**Table 2.**
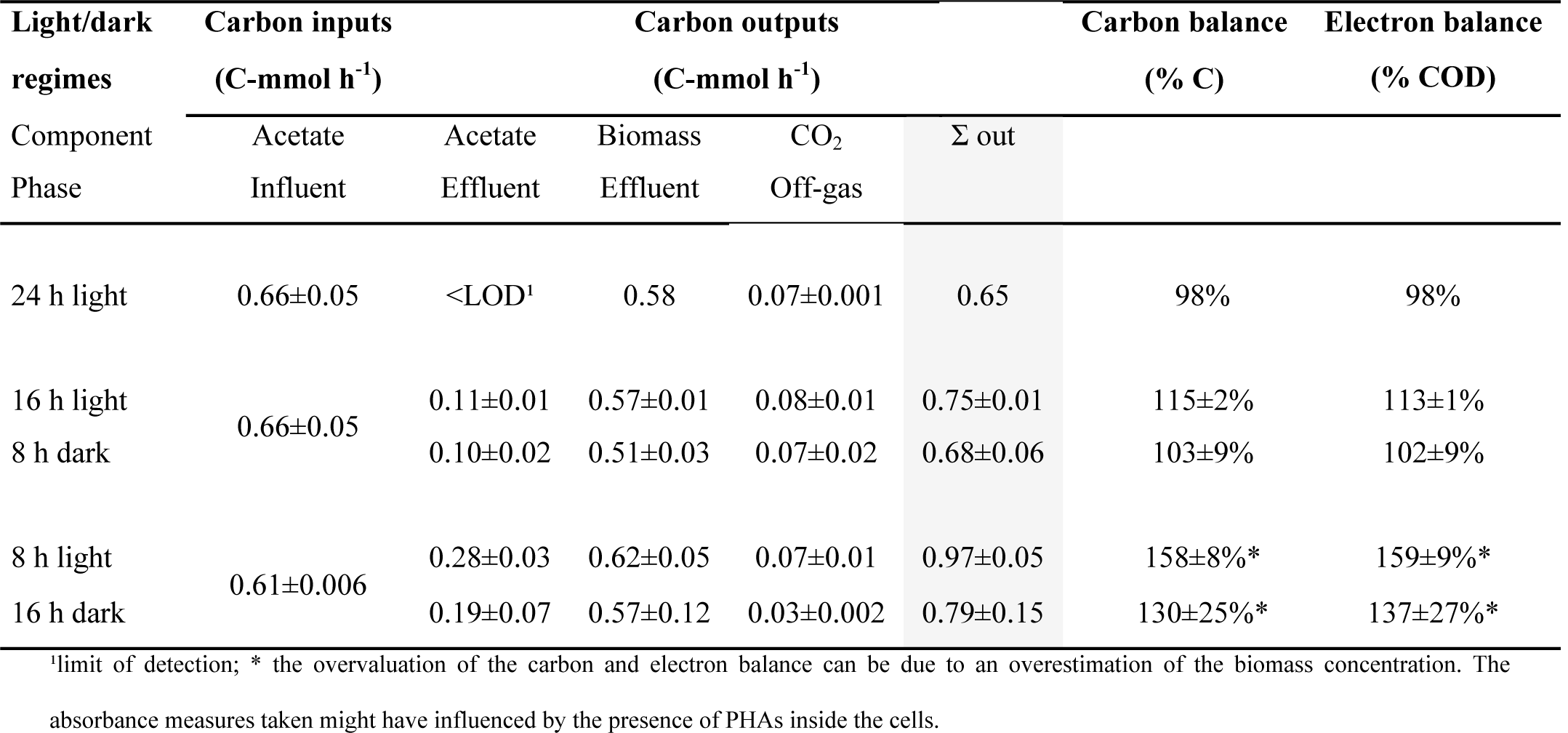
Distribution of the carbon sources and carbon balances under different illumination conditions

Light / dark cycles occur daily in natural environments. Phototrophic organisms have adapted to cope with imbalances in energy supply. Purple bacteria have higher biomass production under light / dark cycles (43,44). H_2_ production is increased under discontinuous illumination, following the biomass trend (45,46).

Here, the light cycles are important not only for the energy that light provides, but most importantly for the time available for the cultures to metabolize the nutrients. The light intensity at the surface of the reactor was 300 W m^-2^. Considering the attenuation due to the biomass concentration and the reactor depth, the minimum calculated light available was 180 W m^-2^ (Figure SI-4). Under any given condition, the available light intensity was not limiting the microbial metabolism since always exceeding the half-saturation coefficient for light, defined at *K*_*i*_ =10 W m^-2^ (39). Purple bacteria adapt the number and type of photosynthetic unit components based on the different light intensities (47), but Imam et al. (48) reported that light may be saturating for PNSB growth already at 100 W m^-2^. We can therefore assume that the irradiance intensity used in our study during the light periods did not deeply affect the metabolic state of the cells.

The continuous cultivation system was initially set-up based on the stoichiometric and kinetic parameters derived from batch experiments. In the continuously illuminated chemostat, the biomass reached a steady-state, acetate was limiting (i.e., low concentrations), and the biomass growth rate equaled to the dilution rate (0.04 h^-1^). Once the light/dark cycles were applied, a disturbance of the steady-state was immediately observed. Under long (16 h) illumination, the biomass consumed the acetate for growth and after an initial increase of the growth rate the steady-state conditions were restored for the remaining two thirds of the light cycles (with a specific growth rate again equal to the dilution rate). Longer (16 h) dark periods led to an increased accumulation of acetate in the bulk liquid (twice as high as in the 8 h dark), since acetate was fed twice longer and there was no consumption in the dark. Due to the shorter light (8 h light) periods, acetate was not fully consumed at the end of the light phases. Therefore, acetate was no more the growth rate limiting compound and the light period was too short to reach a steady state. The growth rate in the light phase exceeded the value of the imposed dilution rate, leading to a transient accumulation of biomass (Figure 2B). The modelled biomass dynamics agreed with the measurements, fitting quantitatively the values for the 16 h light experiment, where the exact steady state values were obtained. When using the same model parameters for the short (8 h) illumination experiment, the calculated amount of biomass formed during the light periods corresponded well to the values measured only for the second and later cycles, while for the first cycle the model underestimated the biomass formed. The disagreement may be related to the carbon imbalance reported in Table 3. Still, given longer illumination in the last model cycle would allow the biomass to reach exactly the same steady state as in the longer light period experiment (Figure SI-3) after about 10 h light and acetate attained the minimum.

The average acetate consumption rates under 16 h light were 0.08 C-mol h^-1^ L^-1^ (Figure SI-5) and under 8 h light were 0.15 C-mol h^-1^ L^-1^ (Figure SI-6). Initially, the biomass grew at the maximum substrate uptake rate. If the light phase was too short (as in the 8 h light periods), acetate was not fully consumed (2.5 ± 1.1 C-mmol L^-1^) and the biomass grew at its maximum growth rate.

### PHA synthesis patterns reflect the metabolic state of the cells

PHAs can be formed under dynamic conditions or because of imbalance in the degree of reduction between biomass and carbon source. PHAs constitute carbon and energy stocks (49,50). In chemostat conditions, carbon was continuously fed to the cultures and cells did not store PHAs. PHA synthesis in purple bacteria has primarily been reported under nitrogen limitation, as an intrinsic mechanism for the redistribution of carbon excess. PHA accumulation is conventionally reported under growth-limiting conditions (51,52) when the carbon sources are available, but the nutrients (such as N, P or S) to produce cellular components are limited resulting in acetyl-CoA accumulation in the cells.

During the anabolic reactions, *Rhodopseudomonas* incorporates acetate in the form of acetyl-CoA and releases at the same time CoA. In highly active cells, the levels of acetyl-CoA are low, but the levels of CoA are high. In contrast, in non-growing cells, acetyl-CoA is not utilized in anabolic processes and accumulates in the cells (53). A high acetyl-CoA to CoA ratio and the presence of NADPH are required for the initiation of the PHA pathway (54).

The ratio between carbon sources and other nutrients is important to maintain the balance between substrate uptake rate and growth rate. The inflow was defined based on the biomass composition of purple bacteria (38), with a C:N ratio of 5.68:1 mol/mol was used in the medium. Nitrogen was provided in excess preventing N-limitation imbalance. Consequently, under continuous illumination, no production of PHAs was detected.

After a period of darkness, the biomass was subjected to an excess of light and nutrients (acetate, ammonium, phosphate) resulting in an increased substrate uptake rate. For the first 2 h after switching on the 16-h light phase, the growth rate was close to the previously measured maximal growth rate μ_max_ of 0.15 h^-1^. After 3 h, when acetate was no more in excess, the growth rate stabilized again at the dilution rate value of 0.04 h^-1^. When shorter (8 h) light periods were applied, the initial growth rate was ∼0.1 h^-1^. Under both conditions, cells exhibited growth rates close to μ_max_. This resulted in the production of reducing power, which, along with the NADH produced in the photosynthetic processes, became in excess and had to be reallocated. As reported by Kanno et al. (55), the photosynthetic units are not disassembled in the dark, even under starvation. These are readily available, once the light conditions are restored, to produce ATP and NADH for the biosynthetic processes. This, linked to the prompt availability of the enzymes for PHAs formation that are constitutively expressed (56), leads to the immediate production of PHAs. Under 16 h light, a peak of fluorescent counts was observed 2 h after the light switch, but it decreased to baseline levels (steady state) once the growth rate stabilized again at 0.04 h^-1^. The fluorescent counts decreased and no H_2_ production was observed. Cells reached a maximal capacity of substrate uptake, though not coupled to a maximal growth rate, similarly to what is described in (57).

Under 8 h light / 16 h dark cycles, acetate was not completely consumed during illumination and increasingly accumulated during the dark phases, resulting in a further imbalance in the redox state of the cells. The redox imbalance generated by the continuous presence of acetate, rather than the illumination conditions alone, led to a more constant production of PHAs. The cells were highly active, with a growth rate close to the maximal growth rate, leading to a low availability of CoA that resulted in PHAs formation. Only when the PHAs pool got saturated the NADPH pool was further increased, and H_2_ production became possible.

### H_2_ production is a secondary pathway of electron dissipation

The H_2_ production rates here reported are low compared to other studies that have exposed PNSB to similar light intensities (48). However, our experimental conditions were not designed to stimulate H_2_ production. H_2_ can be produced either via the nitrogenase system, either via the ferredoxin-hydrogenase system. Ammonium is known to inhibit N_2_ fixation in photosynthetic bacteria. It also effectively prevents photoproduction of H_2_, due to inhibition and inactivation of nitrogenase (18). The presence of ammonium in non-limiting amounts in the medium indicated that potentially H_2_ production was driven by the hydrogenase rather than by the nitrogenase. The molar C:N ratio (5.7) was 7 times lower than in most other studies on H_2_ production (average C:N ratio 40) (58). Nonetheless, the H_2_ production rate per gram biomass during the 8 h light phases was around 11 times higher than in all other conditions (0.156 *vs* 0.014 mmol h^-1^ g^-1^ DW), indicating that H_2_ production acts as further electron dissipation pathway. A similar H_2_ production pattern under alternate irradiation in PNSB cultures has been found (59), with H_2_ production only during the light phases. H_2_ production was not integrated in the numerical model, since H_2_ did not contribute significantly to the stoichiometric balance and kinetics.

The absence of relevant H_2_ production over 16 h light (0.004 ± 0.0004 mmol h^-1^) and its low but constant production during 8 h light (0.025 ± 0.009 mmol h^-1^) indicates that the H_2_ sink did not play a major role in the electron redistribution patterns.

Overall, PNSB are one of the most versatile guilds of microorganisms. They present consistent differences in their metabolisms, interspecies and intraspecies. We presented a comprehensive explanation of the underlying mechanisms of electron allocation using quantitative biotechnology and metabolic modelling. The numerical model represented well the biomass and nutrient dynamics during the light/dark cycles of different durations. It allowed for determining stoichiometric and kinetic parameters of the *Rhodopseudomonas*. It can be concluded that:

1. Under light-saturating conditions and continuous-flow reactor regime, durations of light and dark phases in a diel cycle set the availability of substrate and the achievement of a steady state. Longer dark phases result in an excess of substrate available in the light phase for biomass growth. Longer light phases lead to substrate limitation and steady conditions.
2. Even in actively growing cells, carbon allocation is in place, namely toward PHA production.
3. In growing cells, H_2_ production during illumination is a minor electron sink and secondary to PHA production.

## Material and Methods

### Strain

The *Rhodopseudomonas* strain was isolated with dilution series in agar from an in-house PNSB enrichment culture designed for nutrient removal from synthetic wastewater (30). The isolate was characterized by full-length 16S rRNA gene sequencing. The genomic DNA was extracted using UltraClean® Microbial DNA Isolation Kit (MO BIO Laboratories, Inc., USA), following manufacturer’s instructions. The full 16S rRNA gene was amplified with the primers forward U515 (5’-GTGYCAGCMGCCGCGGTA-3’) and reverse U1071 (5’-GARCTGRCGRCRRCCATGCA-3’) (31), and sequenced for phylogenetic identification using a Sanger sequencing (Baseclear, NL). The sequence was aligned over the Blast database (32), and resulted in a 97.2% identity with *Rhodopseudomonas palustris*.

### Medium

The inflow medium was adapted from Cerruti et al. (30). It consisted of (per liter): 0.914 g CH_3_COONa·3H_2_O (13.5 C-mmol L^-1^ or 54 mmol electrons L^-1^ when expressed via degree of reduction and 432 mg COD L^-1^ when expressed as chemical oxygen demand), 0.229 g NH_4_Cl (*i.e*., 4.281 mmol N L^-1^ or 60 mg N-NH_4_^+^ L^-1^), 0.014 g KH_2_PO_4_ and 0.021 g K_2_HPO_4_ (*i.e*., 0.223 mmol P L^-1^ or 7 mg P L^-1^), 0.200 g MgSO_4_·7H_2_O, 0.200 g NaCl, 0.050 g CaCl_2_·2H_2_O, 1 mL vitamins solution, 1 mL trace element solution, and 4.7 g 4-(2-hydroxyethyl)-1-piperazineethanesulfonic acid (HEPES) used as pH buffer.

The vitamin solution was composed of (per liter): 200 mg thiamine–HCl, 500 mg niacin, 300 mg ρ-amino-benzoic acid, 100 mg pyridoxine–HCl, 50 mg biotin and 50 mg vitamin B12.

The trace element solution was composed of (per liter): 1100 mg Na EDTA·2H_2_O, 2000 mg FeCl_3_·6H2O, 100 mg ZnCl_2_, 64 mg MnSO_4_·H_2_O, 100 mg H_3_BO_3_, 100 mg CoCl_2_·6H_2_O, 24 mg Na_2_MoO_4_·2H_2_O, 16 mg CuSO_4_·5H_2_O, 10 mg NiCl_2_·6H_2_O and 5 mg NaSeO_3_.

The medium components were sterilized though autoclavation or filtration with 0.22 μm filters (Whatman, USA) (acetate solution, trace elements and vitamins).

### Reactor setup

Continuous cultures were run to evaluate the influence of diel cycles on the physiology of PNSB. A 1.5-L continuous-flow stirred-tank reactor with 1.2-L working volume was connected to a programmable logic controller (In-Control and Power unit, Applikon, NL) and operated under stirring of 350 rpm, pH 7.0 ± 0.5, temperature of 30±1 °C during illumination phases and 20°C during dark phases. Argon gas (Linde, NL, >99% purity) was sparged continuously in the bulk liquid phase at 120 mL min^-1^ to maintain anaerobic conditions. The continuous-flow rate was set at 0.048 L h^-1^. It corresponded to a dilution rate of 0.04 h^-1^ chosen based on the growth rate of the strain previously measured at 0.11 h^-1^ (data not shown). The biomass was maintained at a low concentration of 0.26 ± 0.05 g DW L^-1^ to minimize light shading effects.

The reactor was placed in a shaded hood to tightly control the irradiation patterns. Two halogen floodlight lamps (Handson, NL) were positioned at opposite sides of the reactor diameter. The incident white light spectrum was filtered for infrared (IR) light (λ > 700 nm) using two filter sheets of 70 × 70 cm (Black Perspex 962, Plasticstockist, UK). The IR light intensity measured at the reactor surface with a pyranometer (CMP3; Kipp & Zonen, NL) was 300 W m^-2^. An automatic device was used to switch on/off the light at required time sets. Three light conditions were tested: 1) continuous illumination (*i.e*., 24 h light), 2) 16 h light and 8 h dark cycles, and 3) 8 h light and 16 h dark cycles.

### Analytical methods

The biomass concentration was measured by absorbance at 660 nm (A_660_) using a spectrophotometer (Biochrom, Libra S11, USA). A calibration curve was established to correlate A_660_ to dry weight (DW) concentration: c(g DW L^-1^) = 0.64 A_660_ – 0.06. Biomass dry weight was measured taking samples from the liquid phase, filtering them using 0.45 μm filters (Whatman, USA) and storing them in a 70 °C stove for 72 h (adapted from Lip et al. 2020).

Acetate was measured with a high-performance liquid chromatograph (HPLC) (Waters, 2707, NL) equipped with an Aminex HPX-87H column (BioRad, USA). Samples were eluted using H_3_PO_4_ (1.5 mmol L^-1^, flowrate of 0.6 mL min^-1^, temperature of 60°C) prior to refraction index (Waters 2414) and UV (210 nm, Waters 484) detections.

Ammonium (as N-NH_4_^+^) and orthophosphate (as P-PO_4_^3-^) concentrations were measured with a discrete analyser (Thermoscientific Gallery, NL).

PHA measurements were performed with fluorimetry for high sensitivity on small biomass samples. The low biomass concentration did not allow for traditional extractions of PHA and gas chromatographic measurements of monomers. PHAs were stained in the biological samples with Nile red (CAS n. 7385-67-3, Sigma Aldricht), as in (34). Fluorescence was measured with a microplate reader (1000M pro, Tecan), with excitation at 535 nm and emission at 605 nm. The fluorescent counts were normalized by the g DW of biomass present in analysed samples. The absorbance at 660 nm was measured in each well, and then converted in gDW biomass, to this end.

The CO_2_ and H_2_ in the offgas were measured using a mass spectrometer (Thermofisher, Prima BT Benchtop MS) connected online to the bioreactor. The production rates of these components were calculated using the argon gas inflow rate (120 mL h^-1^).

### Mathematical model

A simple mathematical model was developed to characterize the behavior of the continuous culture of *Rhodopseudomonas* exposed to light/dark cycles. Under light, photoorganoheterotrophic growth was assumed; under dark, completely ceased metabolic activity. The growth stoichiometry with light as energy source followed equation (1), with the biomass composition adapted from (35):

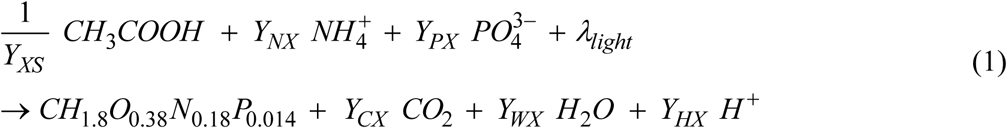

The process was described by a system of balance equations for the relevant materials in the mixed liquor (eqns. (2) (biomass *C*_*X*_, acetic acid *C*_*S*_, ammonium *C*_*N*_, phosphate *C*_*P*_ and carbon dioxide *C*_*C*_).

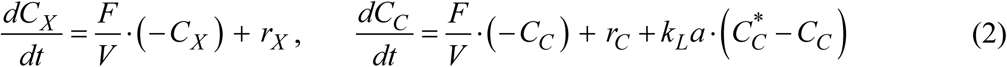

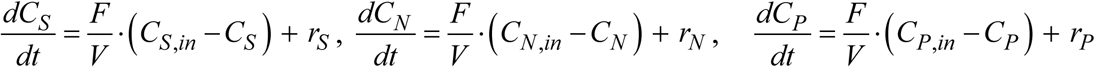

A gas phase balance was integrated for the CO_2_ concentration in the gas *C*_*C,g*_ (mmol/L):

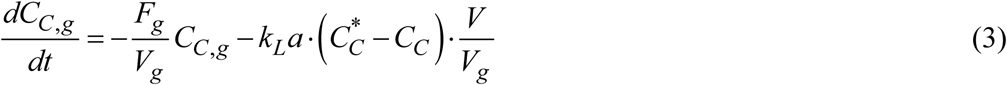

The volumetric growth rate, *r*_*X*_ (C-mmol X L^-1^ h^-1^), was assumed to be limited by the concentrations of multiple chemical compounds and by light intensity *I* (W m^-2^):

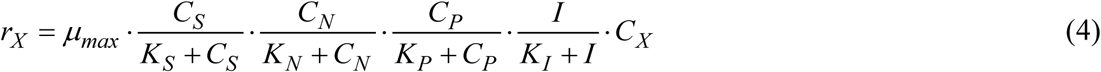

The uptake and production rates of chemical compounds (mmol L^-1^ h^-1^) follow from the reaction stoichiometry and biomass growth rate:

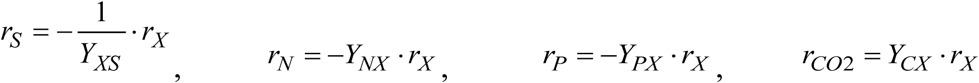

Similarly, the biomass-specific rates were calculated as *q*_*S*_ = *μ* / *Y*_*XS*_, *q*_*N*_ = *Y*_*NX*_ *μ, q*_*P*_ = *Y*_*PX*_ *μ* and *q*_*C*_ = *Y*_*CX*_ *μ*. The yields *Y*_*XS*_, *Y*_*CX*_ and max. specific growth rate μ_max_ were determined by fitting the measurements from the 16h light / 8h dark experiment with this model. Additional estimations of the μ_max_ based on batches and chemostat experiments under continuous illumination and light / dark patterns resulted in μ_max_ values of 0.14 ±0.05. The yields *Y*_*NX*_ and *Y*_*PX*_ (and consequently the N and P biomass composition *nN* and *nP*, respectively) were determined from the 8h light / 16h dark experiment, for which we were able to record better quality data.

The liquid flow rate *F* (L h^-1^) and volume *V* (L), the gas flow rate *F*_*g*_ (L h^-1^) and volume *V*_*g*_ (L), as well as the concentrations of chemical compounds in the influent *C*_*in*_ (mmol L^-1^) were all fixed in the experiments. The volumetric mass transfer coefficient *k*_*L*_*a* (h^-1^) was determined from the correlation with the power input, liquid volume, and gas velocity (see SI-1). The dissolved CO_2_ concentration, 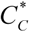 (mmol L^-1^), was calculated function of gas concentration *C* _*C, g*_, Henry coefficient *H*_*C*_ (mol L^-1^ atm^-1^) and respective temperature (20°C in dark and 30°C in light) and pressure (1 atm).

Two light sources with intensity *I*_*0*_ were placed at opposite sides next to the reactor. The light intensity decrease away from the reactor wall was assumed to follow the Beer-Lambert attenuation law. Therefore, by summing the light intensities coming from both sides one obtains the total light intensity *I*_*t*_ at a radial position *x* (m) in the reactor:

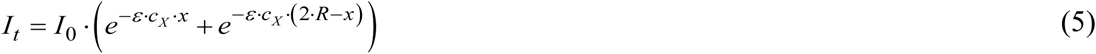

with *R* (m) the reactor radius, ε (m^2^ g^-1^) the biomass-specific light attenuation coefficient and *C*_*X*_ (g m^-3^) the biomass concentration. We considered that due to liquid mixing the cells will be exposed to an average light intensity *I* (W m^-2^) computed by integrating eq. (5) over the reactor diameter:

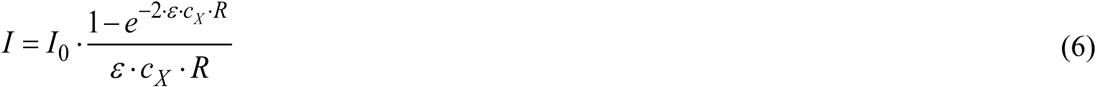

The model was solved in MATLAB (R2018b, Mathworks, Natick, MA, www.mathworks.com) using a stiff solver for the ordinary differential equations system (2) and (3). The initial concentrations where taken to coincide with the measurements. The parameter estimation was performed by a constrained optimization routine minimizing the sum of squares of relative errors between model and experimental data. The fixed model parameters and the fitted parameters are listed in Table 1. A more extensive description of the parameter used is available in SI-1.

## Acknowledgements

This study was financed by the tenure-track start-up grant of the Department of Biotechnology of the Faculty of Applied Sciences of the TU Delft (David Weissbrodt, PI). We acknowledge the Onassis Foundation for the financial support in the scholarship of Viktor Chasna. We also thank Dirk Geerts and Rob Kerste for technical assistance with the reactor infrastructure and fermentation facility, Katie Thorp for the isolation of the culture and Gijs Kuenen for the valuable discussions.

## Preprint

This manuscript will be deposited as pre-print in bioRxiv.

## Supplementary information

Appendix 1: Estimation of the kLa of the process

Appendixes 2: Mathematical model simulations

- Fig. A-1: Data and simulation of the 8 h light / 16 h dark cycles with extended light phase.
- Fig. A-2: Model results for the light intensity inside the reactor
- Fig. A-3: Model results for production and uptake rate in the 16 h light / 8 h dark cycles
- Fig. A-4: Model results for production and uptake rate in the 8 h light / 16 h dark cycles.

## Tables

**Table 1.** Model parameters. The maximum biomass-specific growth rate and yields were fitted to experimental data, while the other parameters were measured during experiments, calculated or obtained from literature.

| Parameter name | Symbol | Value | Units | Source |
| --- | --- | --- | --- | --- |
| Max. biomass-specific growth rate | $\mu_{\max}$ | 0.15 | $\text{C-mol X} \cdot \text{h}^{-1} \cdot \text{C-mol}^{-1} \text{X}$ | Fitted |
| Biomass yield | $Y_{XS}$ | 0.88 | $\text{C-mol X} \cdot \text{C-mol}^{-1} \text{S}$ | Fitted |
| Ammonium yield | $Y_{NX}$ | 0.18 | $\text{mol} \cdot \text{C-mol}^{-1} \text{X}$ | Fitted |
| Phosphate yield | $Y_{PX}$ | 0.009 | $\text{mol} \cdot \text{C-mol}^{-1} \text{X}$ | Fitted |
| CO<sub>2</sub> yield | $Y_{CX}$ | 0.142 | $\text{mol} \cdot \text{C-mol}^{-1} \text{X}$ | Fitted |
| Reactor radius | $R$ | 0.055 | m | Measured |
| Liquid volume | $V$ | 1.2 | L | Measured |
| Liquid flow rate | $F$ | 0.048 | $\text{L} \cdot \text{h}^{-1}$ | Measured |
| Dilution rate | $D$ | 0.04 | $\text{h}^{-1}$ | $F/V$ |
| Gas volume | $V_g$ | 0.3 | L | Measured |
| Gas flow rate | $F_g$ | 7.14 | $\text{L} \cdot \text{h}^{-1}$ | Measured |
| Inflow acetate concentration | $C_{S,in}$ | 7 | mM | Measured |
| Inflow ammonium concentration | $C_{N,in}$ | 4.28 | mM | Measured |
| Inflow phosphate concentration | $C_{P,in}$ | 0.22 | mM | Measured |
| Light intensity at reactor walls | $I_0$ | 300 | $\text{W} \cdot \text{m}^{-2}$ | Measured |
| Light extinction coefficient per biomass concentration | $\varepsilon$ | 0.1 | $\text{m}^2 \cdot \text{g}^{-1}$ | (17) |
| Half-saturation coefficient acetate | $K_S$ | 0.1 | mM | Lower than in (36) (0.3 mM) |
| Half-saturation coefficient ammonium | $K_N$ | 0.001 | mM | (37) |
| Half-saturation coefficient phosphate | $K_P$ | 0.003 | mM | (38) |
| Half-saturation coefficient light | $K_I$ | 10 | $\text{W} \cdot \text{m}^{-2}$ | (39) |
| CO<sub>2</sub> mass transfer coefficient | $k_{La}$ | 32 | $\text{h}^{-1}$ | Calculated (SI-1) |
| Henry coefficient CO<sub>2</sub> | $H_C$ | 0.03 (at 30°C)<br>0.04 (at 20°C) | $\text{mol} \cdot \text{L}^{-1} \cdot \text{atm}^{-1}$ | (40) |

**Table 2.**
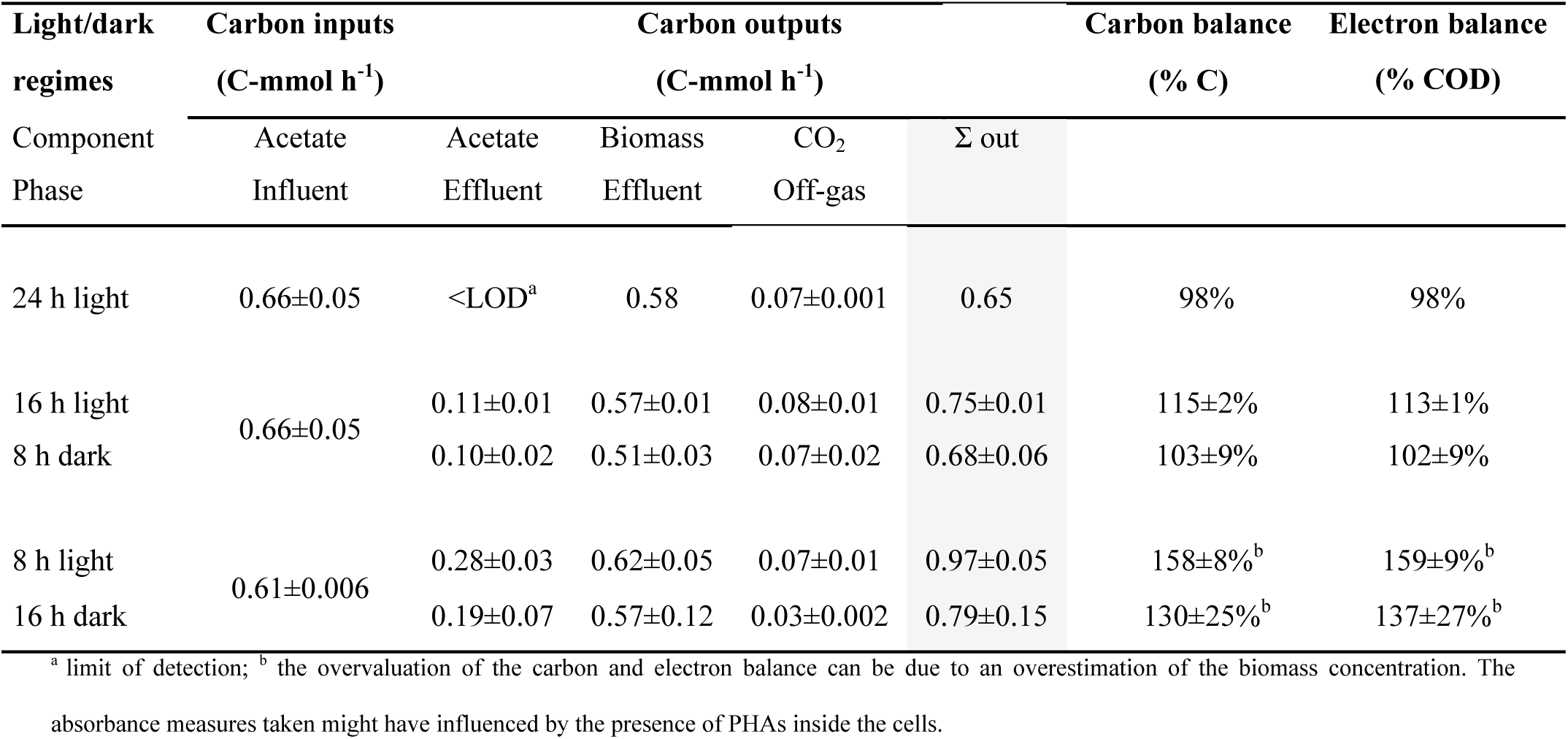
Distribution of the carbon sources and carbon balances in different illumination conditions

## Supplementary information

## Appendix 1: Estimation of the kLa’s of the process

The overall mass transfer coefficients of the process were found in the CO_2_ and O_2_ mass balances. The calculation of the oxygen mass transfer coefficient was performed considering the equation for salt solutions

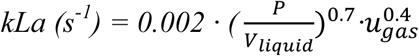

where P is the power input (W), *u*_*gas*_ is the superficial gas velocity (m/s) and *V*_*liquid*_ the liquid volume (m3). The power input was calculated through the following equation (60)

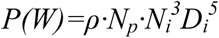

where ρ is the density (kg m^-3^) of the solution, N_p_ is the impeller power number, N_i_ is the impeller rotating speed (s-1) and D_i_ is the impeller diameter(m).

The power number of the impeller is a function of the impeller type and the Reynolds number of the impeller. Reynolds was calculated through the following equation

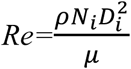

where μ is the viscosity of the solvent (Pa·s). Considering the density (997 kg m^-3^) and viscosity of water in 25 ^°^C (0.00089 Pa·s) the Reynolds impeller was slightly higher than 10^5^. For that Reynolds and for a 6-blade Rushton turbine impeller the power number is around 5∼6 (60). For the calculations, a power number of 6 was assumed. The diameter of the impeller was around 4 cm and the rotating speed 350 rpm (5.83 s^-1^). Using those values the power input of the impeller was found equal to 0.101 W.

The superficial gas velocity was calculated by dividing the gas flowrate (7.14 L h^-1^) with the surface area (π R^2^). The reactor radius R was 5.5 cm. After dividing and correcting for the units the calculated superficial gas velocity was 0.000209 m s^-1^. Using the calculated values of the superficial gas velocity, power input and liquid volume (1.2 ·10^−3^ m^3^) the k_L_a of oxygen was found equal to 33.52 (h^-1^).

The k_L_a of carbon dioxide was found equal to 32 (h^-1^) through the following equation (61). The diffusion coefficient used are 1.92·10^−5^ cm^2^ s^-1^ (62)

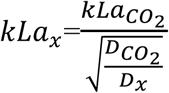

## Appendixes 2 to 5: Mathematical model simulations

**Figure A-2.**
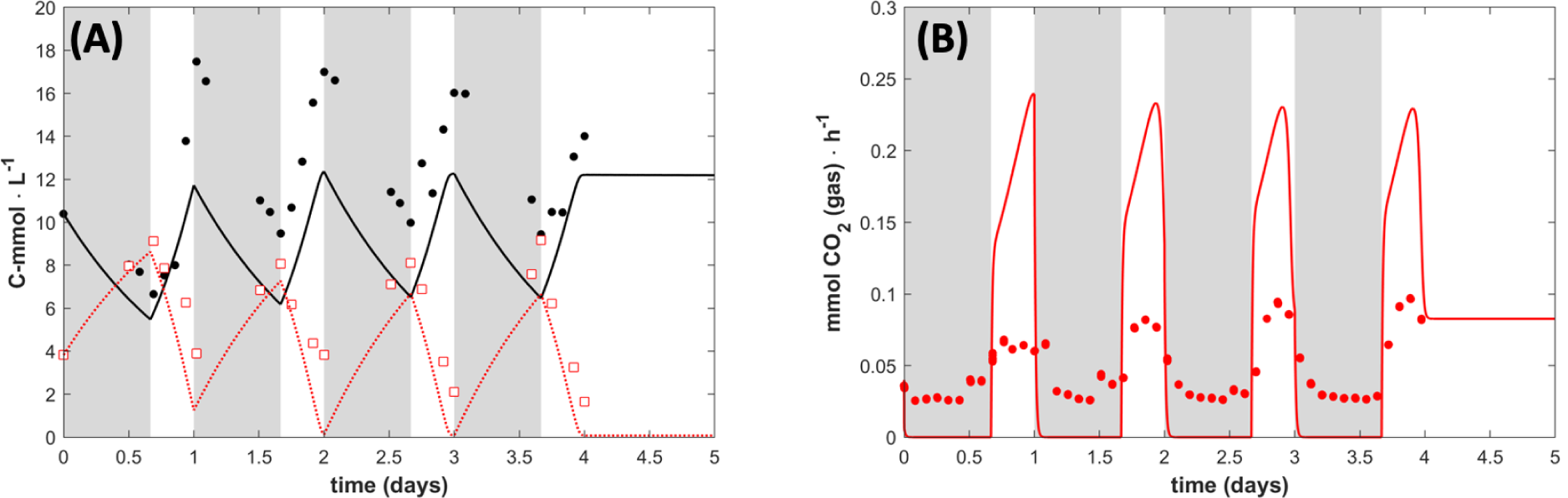
Data and simulation with an additional extended light phase after the 8 h light 16 h dark cycles. (A) Biomass (black circles) and acetate (red open squares) measured concentrations, with lines being the model results. (B) Measured (red circles) and model prediction (line) of the CO_2_ production rate A steady state is reached once the concentration of acetate is at its minimum.

**Figure A-3.**
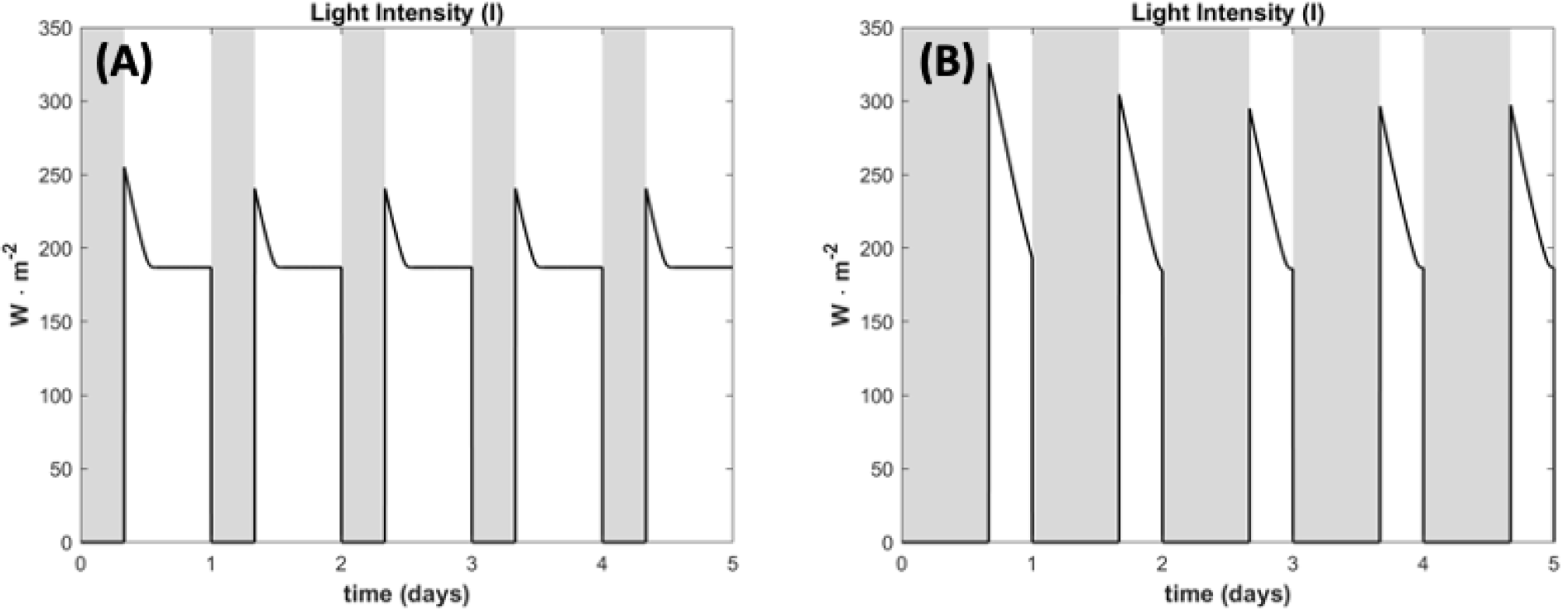
Model results for the light intensity inside the reactor. (A) 16 h light / 8 h dark cycles. (B) 8 h light / 16 h dark cycles. Under any given condition light became a limiting factor.

**Figure A-4.**
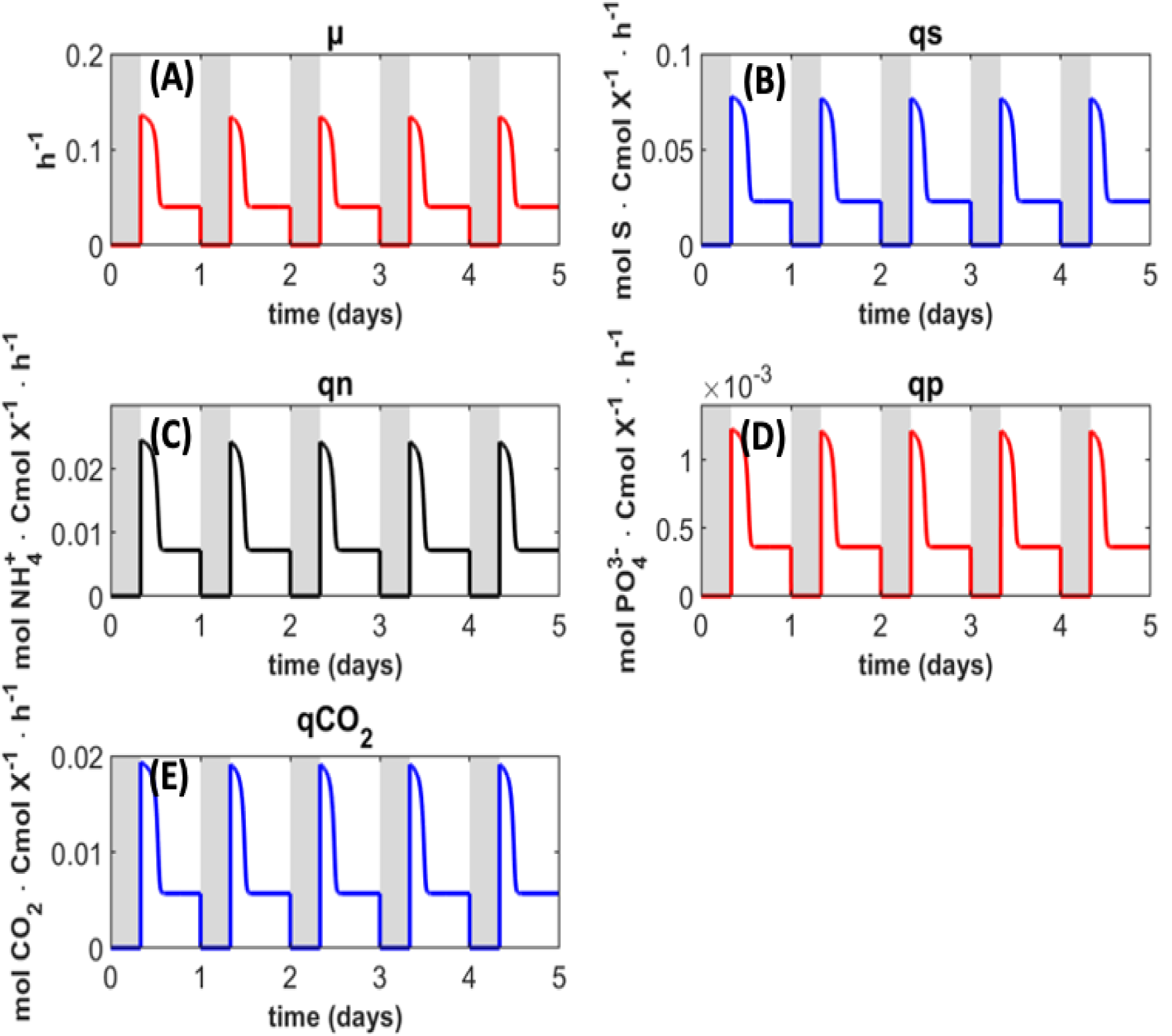
Model results for production and uptake rate in the 16 h light / 8 h dark cycles. (A) specific growth rate. (B specific acetate uptake rate. (C) specific ammonium uptake rate. (D) specific phosphate uptake rate. (E) specific CO_2_ production rate.

**Figure A-5.**
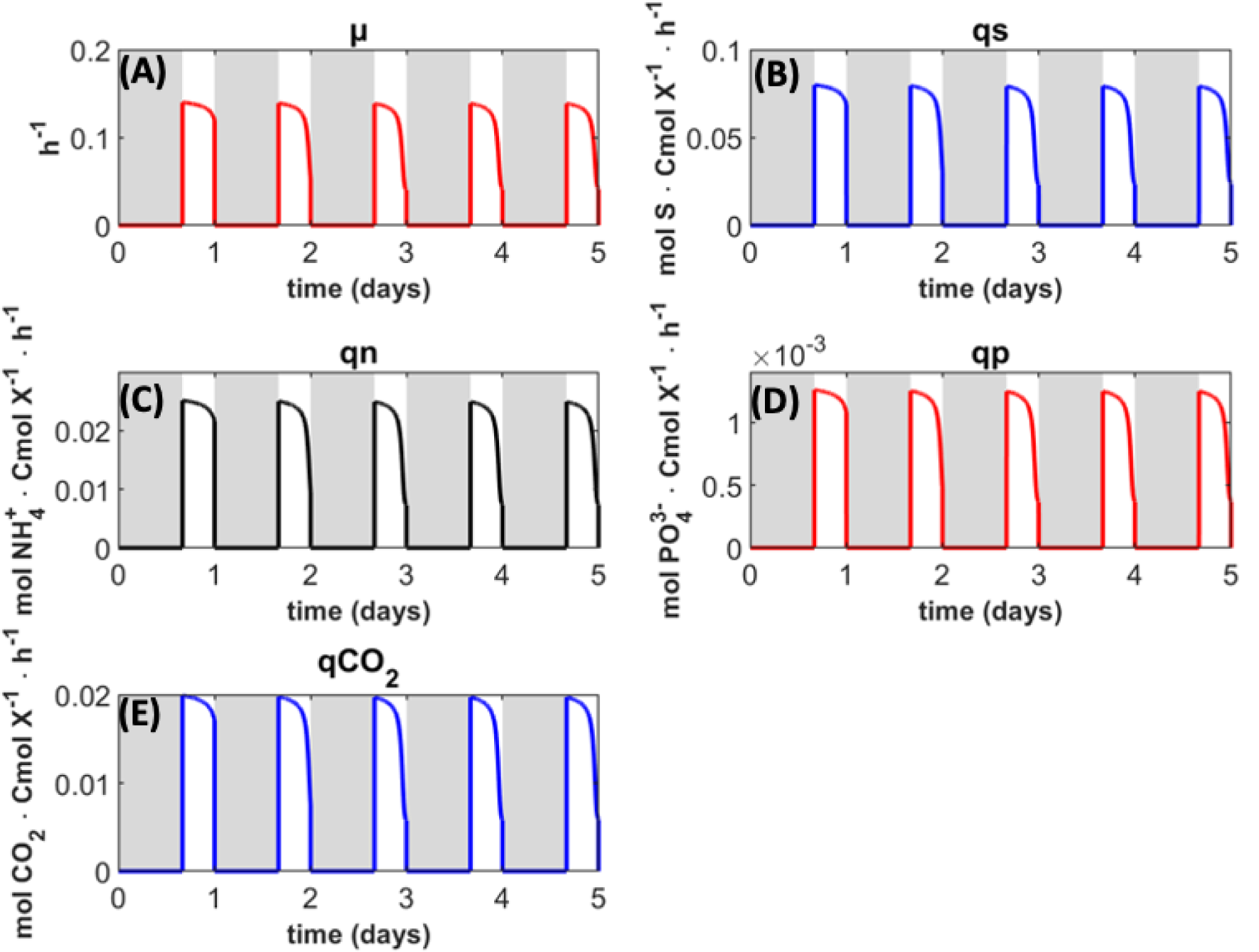
Model results for production and uptake rate in the 8 h light / 16 h dark cycles. (A) specific growth rate. (B specific acetate uptake rate. (C) specific ammonium uptake rate. (D) specific phosphate uptake rate. (E) specific CO_2_ production rate.

